# Undersampling correction methods to control γ-dependence for comparing β-diversity between regions

**DOI:** 10.1101/2021.01.24.427952

**Authors:** Ke Cao, Jens-Christian Svenning, Chuan Yan, Jintun Zhang, Xiangcheng Mi, Keping Ma

## Abstract

Measures of β-diversity are known to be highly constrained by the variation in γ-diversity across regions (i.e., γ-dependence), making it challenging to infer underlying ecological processes. Undersampling correction methods have attempted to estimate the actual β-diversity in order to minimize the effects of γ-dependence arising from the problem of incomplete sampling. However, no study has systematically tested their effectiveness in removing γ-dependence, and examined how well undersampling-corrected β-metrics reflect true β-diversity patterns that respond to ecological gradients. Here, we conduct these tests by comparing two undersampling correction methods with the widely used individual-based null model approach, using both empirical data and simulated communities along a known ecological gradient across a wide range of γ-diversity and sample sizes. We found that undersampling correction methods using diversity accumulation curves were generally more effective than the null model approach in removing γ-dependence. In particular, the undersampling-corrected β-Shannon diversity index was most independent on γ-diversity and was the most reflective of the true β-diversity pattern along the ecological gradient. Moreover, the null model-corrected Jaccard-Chao index removed γ-dependence more effectively than either approach alone. Our validation of undersampling correction methods as effective tools for accommodating γ-dependence greatly facilitates the comparison of β-diversity across regions.

## INTRODUCTION

β-diversity is defined as the difference in species composition across space (Anderson *et al.* 2011). Importantly, analyzing differences in β-diversity across regions allows ecologists to test hypotheses regarding the processes driving patterns of biodiversity (Anderson *et al.* 2011; Mori *et al.* 2018). Recently, researchers have become increasingly aware of the influence of variation in γ-diversity (i.e., the total species richness in a region) on measures of β-diversity (i.e., γ-dependence) (Kraft *et al.* 2011; Myers and LaManna 2016). Typically, the more species that are present in a community, the larger the sample size needed to adequately describe the diversity of the community (Colwell and Coddington 1994; Chao and Jost 2012). As a result, measures of β-diversity across species-poor communities may be relatively free of γ-dependence, since small samples will capture most of the true composition. In contrast, small samples from species-rich communities will likely encompass only a tiny fraction of the true composition, leading to severe undersampling and inflated β-diversity (Condit *et al.* 2005; Tuomisto and Ruokolainen 2012). Therefore, γ-dependence is expected to be ubiquitous and particularly problematic when comparing β-diversity among high- and low-diversity regions (Kraft *et al.* 2011; Tuomisto and Ruokolainen 2012).

Properly accounting for the γ-dependence of β-diversity metrics is important, since γ-dependence can lead to spurious interpretations regarding the ecological mechanisms driving community assembly and dynamics (Myers and LaManna 2016). An individual-based randomization null model approach has been widely used for the correction of γ-dependence (Chase and Myers 2011; Kraft *et al.* 2011; Xu *et al.* 2015; Myers and LaManna 2016). However, its effectiveness in removing γ-dependence has been questioned (Qian *et al.* 2013; Bennett and Gilbert 2016; Ulrich *et al.* 2017). Alternatively, another approach for minimizing the effects of γ-dependence is to simply use β-diversity metrics that explicitly account for undersampling (Colwell and Coddington 1994; Chao and Jost 2012; Marcon *et al.* 2012; Tuomisto and Ruokolainen 2012). For example, estimators have been developed to adjust the Jaccard and Sørensen indices based on the degree of undersampling in the data (Chao *et al.* 2005). Species accumulation curves have been applied to rarefy and extrapolate species richness with respect to sample size (Colwell *et al.*, 2012); these methods have been recently extended to diversity accumulation curves to obtain asymptotic estimations of the real β-diversity (Chao *et al.* 2013; 2014).

There is a long history documenting the effects of undersampling on β-diversity (Wolda 1981; Colwell and Coddington 1994; Tuomisto and Ruokolainen 2012; Beck *et al.* 2013), which revealed the necessity for combining undersampling correction methods to facilitate the comparison of β-diversity across regions. However, no study has examined whether β-diversity metrics incorporated with specific undersampling correction methods could effectively remove γ-dependence. While previous studies have examined the robustness of β-metrics to γ-dependence by testing whether metrics can identify simple unstructured communities randomly sampled from species pools of different sizes (e.g. Kraft *et al.* 2011; Bennett and Gilbert 2016; Ulrich *et al.* 2017), this method may not reflect natural communities that are likely structured by ecological gradients. Therefore, a rigorous assessment of the ability of β-diversity measures to reveal true β-diversity patterns in response to ecological gradients is needed.

We compared the effectiveness—in terms of the magnitude of independence on γ-diversity and sample size—of two undersampling correction methods to the null model approach in conjunction with two major classes of commonly used β-diversity metrics. To do this, we used both Gentry’s global forest dataset and simulated metacommunities with variable degrees of stochastic and deterministic responses to a known ecological gradient, as well as a wide range of sample sizes and γ-diversities. Using these data, we asked the following questions: (1) Are β-metrics that incorporate undersampling correction methods able to effectively remove γ-dependence compared with similar uncorrected β-metrics? (2) Do β-metrics with undersampling correction methods reflect real β-diversity patterns caused by underlying ecological processes? and (3) Do undersampling correction methods outperform null model approaches in removing γ-dependence and reflecting ecological gradients? We expect that results from this study will lead to useful insights regarding the most appropriate β-diversity metrics to use across different communities around the globe.

## METHODS

### Abundance-basedβ-metrics

We selected five commonly used abundance-based β-metrics, representing two general approaches to measure β-diversity (Anderson *et al.* 2011): classical metrics calculated using α-diversity and γ-diversity, and multivariate metrics based on summary statistics of pairwise dissimilarity among samples (Baselga 2010; Legendre and De Caceres 2013). Although these two classes of metrics emphasize different facets of β-diversity, they capture Whittaker’s original measures of β-diversity as variation in species composition along environmental or spatial gradients (Anderson *et al.* 2011; Legendre and De Caceres 2013).

In the two classes of β-metrics, we first excluded β-metrics that were mathematically dependent on γ-diversity (Chao *et al.* 2012), because the information these metrics contain about γ-diversity may result in correlations with β-diversity (Tuomisto 2010; Chao *et al.* 2012; Marcon *et al.* 2012; Legendre and De Caceres 2013). Next, we focused on abundance-based β-diversity metrics, since the γ-dependence of incidence-based β metrics (metrics based on species presence-absence data) have been explored in previous studies (Bennett and Gilbert 2016), and accurate undersampling correction is difficult to obtain based on incidence data alone (Chao *et al.* 2006). Therefore, of the classical metrics, we considered the β-Shannon diversity and the normalized divergence indices (Jost 2007; Marcon *et al.* 2012; Chao and Chiu 2016). These two metrics quantify β-diversity as the effective number of compositionally distinct sampling units, which is equal to the “true β-diversity” defined by Jost (2007) and Tuomisto (2010, see more details in *Appendix 2*). Multivariate metrics have been shown to be more robust to γ-dependence than classical metrics (Condit *et al.* 2005; Bennett and Gilbert 2016; Marion *et al.* 2017), and we were especially interested in comparing β-diversity across multiple communities; thus we chose to examine the widely-used Jaccard, Hellinger, and Bray-Curtis pairwise dissimilarity indices, and transformed pairwise dissimilarity matrices into the total variance of community compositional heterogeneity (Legendre and De Caceres 2013. See details in *Appendix 1*).

For classical metrics, we assayed the effectiveness of undersampling correction methods by comparing the undersampling-corrected β-Shannon diversity (Chao *et al.* 2013; 2014) to the raw normalized divergence index (Chao and Chiu 2016). We conducted this comparison because the β-Shannon diversity and the normalized divergence index are exactly identical (*q* = 1) given true species richness and species abundances (Chao *et al.* 2019). For the multivariate metrics, we compared the total variance of the undersampling-corrected Jaccard-Chao distance matrix (Chao *et al.* 2005) to the total variance of the Hellinger and Bray-Curtis distance matrices.

### Undersampling correction methods and the null model approach

For multivariate metrics, Chao *et al.* (2005) proposed the undersampling-corrected Jaccard index (Jaccard-Chao), which estimates the relative abundances of undetected shared species (see more details in Appendix 2). Classical metrics are calculated based on γ- and α-diversity, and observed diversity at both α- and γ-scales are constrained by undersampling bias (Chao *et al.* 2013; 2014). Chao *et al.* (2013; 2014) extended the species accumulation curve (Colwell *et al.* 2012) to the diversity accumulation curve, which corrects for the γ-dependence of β-Shannon diversity by asymptotically estimating both true α- and γ-Shannon diversity 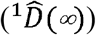) of samples in a region (See details of 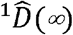 in *Appendix 2*).

As a comparison, we also examined the effectiveness of the randomization null model by calculating all five null model-corrected β-diversity measures (β-deviations) (Chase and Myers 2011; Kraft *et al.* 2011) (see more details of null model approach in Appendix 2).

### Simulated metacommunities and empirical data

We applied a niche-based competition model to create a total of 16200 metacommunities. These metacommunites were produced under 324 simulation scenarios with nine levels of niche strength from neutral to niche-structured, in combination with six levels of γ-diversity (50, 100, 150, 200, 300, and 400 species) and six different sample sizes (50, 100, 150, 200, 250, and 300 individuals per community) (See more details in *Appendix 3*), and each scenario was executed for 50 replicates. We set the metacommunity scale as the γ-scale, and each community as the α-scale.

We also downloaded Gentry’s global forest dataset from SALVIAS (www.salvias.net). The dataset consists of 197 sampling plots distributed from temperate to tropical forests around the world (Phillips and Miller 2002). Within each plot, ten 0.01 ha (2 m × 50 m) subplots were surveyed, where all woody individuals with a diameter of breast height (DBH):2 2.5 cm were identified and recorded. We set the plot scale (i.e., all ten subplots) as the γ-scale, and each subplot as α-scale.

### Statistical analyses

To compare the performance of β-metrics, we regressed the raw and undersampling-corrected β-diversity metrics and their β-deviations against γ-diversity and sample size using multiple linear regression. All variables were standardized before being included in the model. To assess whether β-diversity metrics and their β-deviations were able to distinguish β-diversity created by different simulated niche strength scenarios, we examined the significance of differences in β-diversity among niche-strength scenarios. Finally, we performed an analysis of variance (ANOVA) to analyze the differences of β-diversity among niche strength scenarios, followed by a multiple comparisons based on Tukey’s honestly significant difference (HSD) test (Tukey 1949). All statistical analyses were performed in R, version 3.4.1 (R Core Team 2019). The Shannon diversity index and all undersampling corrections were implemented using the “entropart” package (Marcon and Hérault 2015). The Hellinger, Bray-Curtis, and Jaccard-Chao indices were calculated in “vegan” package (Oksanen *et al.* 2015).

## RESULTS

### The effectiveness of undersampling correction methods in correctingγ-dependence

Metrics for comparing β-diversity among regions are not expected to display systematic changes with γ-diversity and sample size. Using simulated data, we found the corrected β-Shannon diversity to be relatively insensitive to changes in γ-diversity and sample size (Fig. 1a and *Table S1*), as the Jaccard-chao index showed much less dependence (Fig. 1d). In contrast, the normalized divergence, Hellinger, and Bray-Curtis indices were strongly dependent on both (Fig. 1b-c, e and *Table S1*). In general, results of Gentry’s datasets confirmed the results of the simulation experiment (*Table S3*), except that the Jaccard-Chao index had a much stronger γ-dependence than in simulated communities (*Table S1* and *S3*).

**Figure 1.**
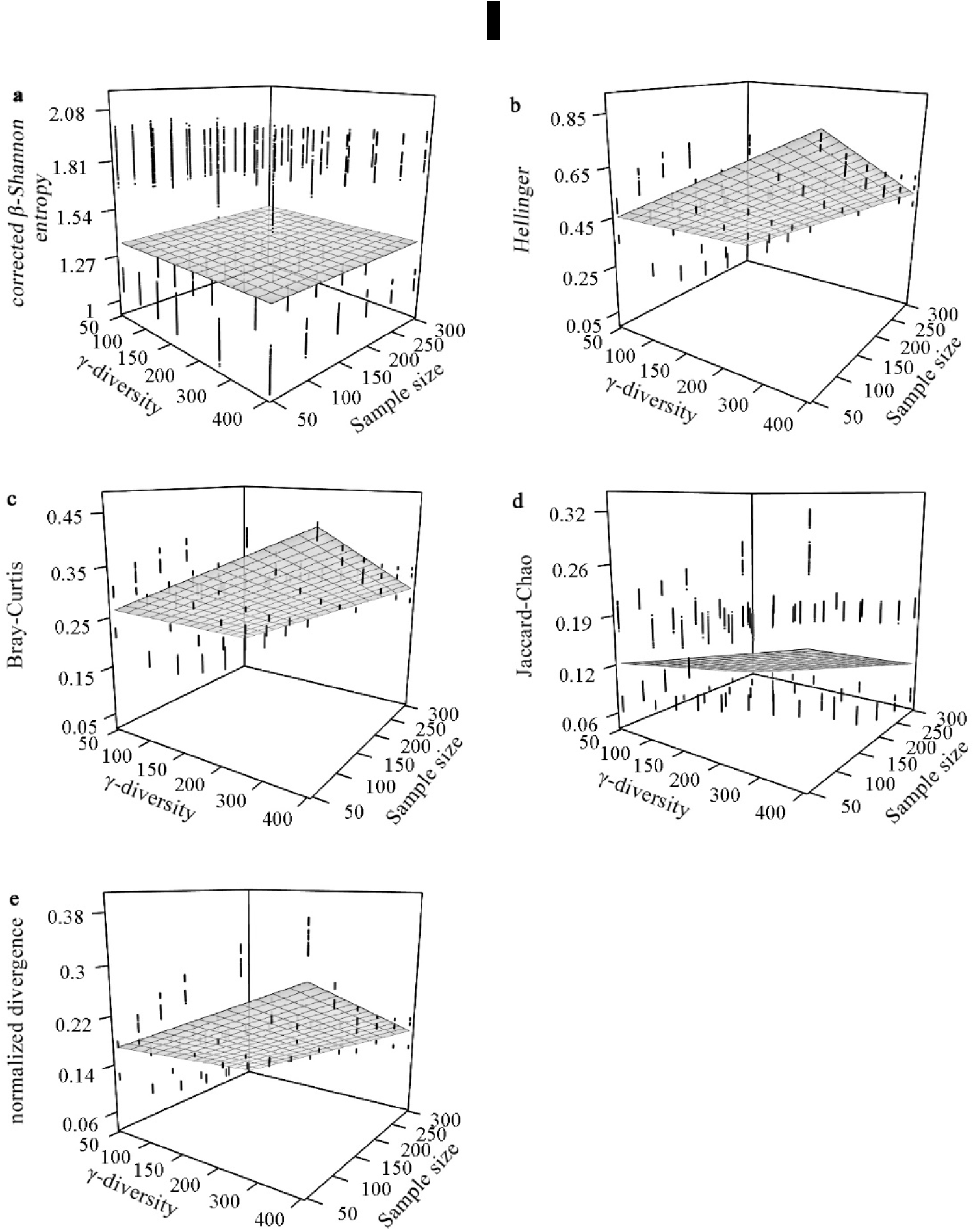
The sensitivity of raw and undersampling corrected β-diversity metrics to γ-diversity and sample size. β-diversity was measured by a) corrected β-Shannon diversity, b) Hellinger, c) Bray-Curtis, d) Jaccard-Chao index, and e) the normalized divergence index. In each panel, the surface of γ-diversity and sample size was fitted using multiple linear regression (detailed model parameters are listed in *Table S1*).

### Variation ofβ-metrics along a niche strength gradient

Metrics for comparing β-diversity among regions should ideally detect the variation of β-diversity across niche strength scenarios regardless of γ-diversity and sample size. Only the corrected β-Shannon diversity clearly distinguished all niche strength scenarios (Fig. 2a). In contrast, the Hellinger, Bray-Curtis, Jaccard-Chao, and normalized divergence indices only roughly detected the differences in β-diversity between strongly niche-structured scenarios and others; they were unable to distinguish between neutrally-structured metacommunities and those with low and moderate niche-strength (Figs. 2b-2e, the left six groups), nor between the three strongest niche-structured communities.

**Figure 2.**
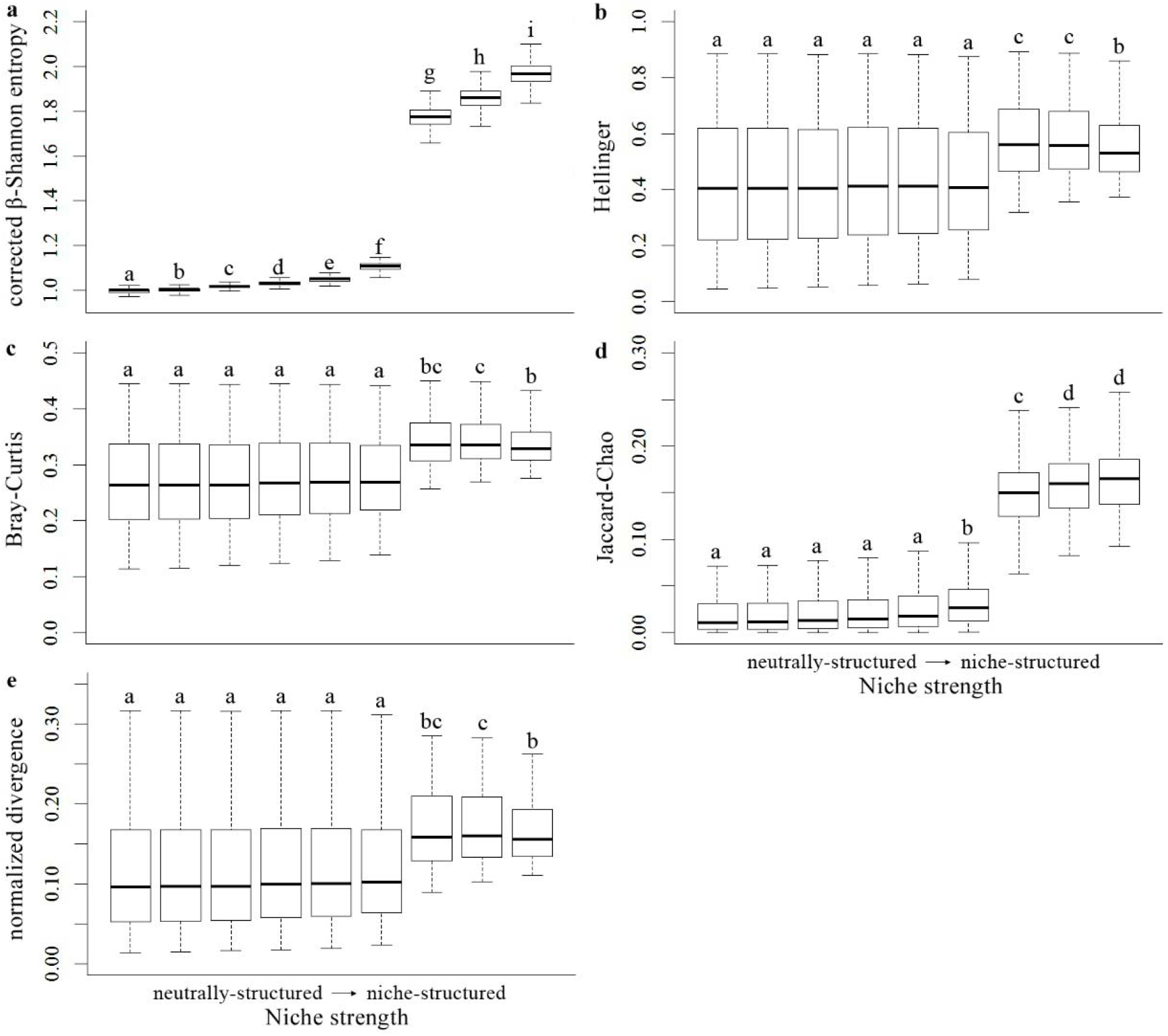
The variation of β-diversity metrics along a niche strength gradient. From left to right, metacommunities varied from stochastically-structured to niche-structured; a) corrected β-Shannon diversity, b) Hellinger, c) Bray-Curtis, d) Jaccard-Chao, and e) normalized divergence indices. Different letters annotated in each panel indicate significantly different mean values of different ecological scenarios.

### The performance ofβ-metrics with a null model approach

Using simulated data, the null model approach generally reduced the dependence of γ-diversity and sample size for all β-metrics tested (*Fig. S1*). The β-deviations of the corrected β-Shannon diversity and Jaccard-Chao index were only slightly sensitive to γ-diversity and sample size (*Fig. S1*, *Table S2*). Compared to the raw normalized divergence, the sensitivities of the β-deviations of the Hellinger and Bray-Curtis indices to γ-diversity and sample size were greatly reduced, albeit non-zero (Fig. *S1*, *Table S2*). Results of empirical data showed very similar results to the simulated data (*Table S4*), except the β-deviation of the normalized divergence index showed greater γ-dependence (*Table S3* and *S4*).

The β-deviations of the raw metrics and the Jaccard-Chao index showed some ability to distinguish between niche strength scenarios; however, these β-deviations were incapable of discriminating between neutrally- and weakly niche-structured metacommunities (*Figs. S2b-S2e*).

## DISCUSSION

We found that the undersampling correction method using the diversity accumulation curve (Chao *et al.* 2014) was most successful at correcting γ-dependence for the β-Shannon diversity and distinguishing β-diversity patterns generated from simulated ecological gradients. As a result, undersampling correction methods using diversity accumulation curves are more promising tools for correcting γ-dependence than current null model approaches. This increased performance is likely because the undersampling correction method approximates the true β-diversity using diversity accumulation curve to separately estimate mathematically independent true α- and γ-diversity (Marcon *et al.* 2012; Chao *et al.* 2014). This avoids the interdependence of β- and γ-diversity (Bennett and Gilbert 2016; Ulrich *et al.* 2017), and the problem of removing a real trend by preserving species abundance distribution in the randomization process of the null model approach (Qian *et al.* 2013; Xu *et al.* 2015). On the other hand, it is worth noting that the effectiveness of specific undersampling correction methods may not be adequate to accurately estimate the true diversity when sample sizes are extremely small (Chao *et al.* 2005; Chao *et al.* 2014). For example, Jaccard-Chao failed in reducing undersampling bias in Gentry’s data caused by severe sampling at both α and γ scales in Gentry’s dataset with an average of 34 individual trees in each sbuplot (Tuomisto and Ruokolainen 2012).

Multivariate metrics were shown to be more robust to γ-dependence than classical measures because the mean α-diversity and the total diversity of each sample pair for pairwise metrics do not increase with the number of sampling units (Bennett and Gilbert 2016; Marion *et al.* 2017). However, our results show that multivariate metrics still suffer from γ-dependence, perhaps because same-sized plots share a smaller fraction of species in higher diversity areas (Condit *et al.* 2005).

β-deviations have become a popular method for comparing β-diversity among regions. However, β-deviations of raw metrics also retained a degree of γ-dependence (Bennett and Gilbert 2016; Ulrich *et al.* 2017), and failed to discern the β-diversity pattern along a known ecological gradient (Bennett and Gilbert 2016). However, the null model approach can be integrated with a wider range of β-metrics and may be more useful than undersampling correction methods when the undersampling of community data is not severe (Chase and Myers 2011; Tucker *et al.* 2016). Moreover, the β-deviation of the Jaccard-Chao index greatly outperformed either approach alone, suggesting a complementary way to combine approaches.

Our results illustrate the importance of testing the robustness of β-metrics to γ-dependence with simulated communities along a known gradient, as it is inadequate to examine the robustness using unstructured communities randomly sampled from species pools of different sizes. In this study, we found that all β-deviations of raw metrics could identify neutrally structured communities (β_*dev*_≈0, *Fig. S2*), but were unable to discern the variation of β-diversity along an ecological gradient.

Taken together, we found that the undersampling correction methods using diversity accumulation curves were more effective at removing γ-dependence than commonly used null model approaches; in particular, the corrected β-Shannon diversity performed best. Meanwhile, the null model-corrected Jaccard-Chao index may exemplify a complementary way to combine a null model approach with a less effective undersampling correction method. These tools allow for comparison of β-diversity along broad biogeographic or disturbance gradients with changing γ-diversity. However, the reliability of different β-metrics at comparing compositional differences between regions depends on other ecosystem characteristics such as species abundance distribution and intraspecific aggregation (Chao and Jost 2012; Beck *et al.* 2013). Other approaches remain to be examined with respect to correcting γ-dependence in future studies. For example, the Simpson dissimilarity index (β_sim_) is supposed to quantify β-diversity due to true spatial turnover of species among sites without the influence of γ-diversity (Baselga 2010).

## ACKNOWLEDGEMENTS

We thank Richard Condit, Chun-Huo Chiu, Tak Fung and Dingliang Xing for their insightful comments on an earlier draft. Funding for this project were provided by the Strategic Priority Research Program of the Chinese Academy of Sciences (XDA19050500) and by the National Natural Science Foundation of China (NSFC 31770478). JCS considers this work a contribution to his VILLUM Investigator project “Biodiversity Dynamics in a Changing World” funded by VILLUM FONDEN (grant 16549).

## Data Availability

The data and associated R code supporting the findings of this study have been uploaded as part of the electronic supplementary material.

---

**Table S1.** The sensitivity of different β-metrics to γ-diversity and sample size based on simulated data. The standardized coefficients are standardized regression coefficients of multiple linear regression models.

| β-metrics | explanatory variables | Standardized coefficients | Standard Error | t value | P(>\|t\|) | R-square | AIC |
| --- | --- | --- | --- | --- | --- | --- | --- |
| **corrected** | γ-diversity | 0.017 | 0.0079 | 2.17 | 0.030 | | |
| **β-Shannon diversity** | sample size | 0.012 | 0.0079 | 1.49 | 0.14 | 0.00043 | 45973.65 |
| **Hellinger** | γ-diversity | 0.67 | 0.0036 | 184.3 | <0.001 | 0.79 | 20779.97 |
|  | sample size | -0.59 | 0.0036 | -163.1 | <0.001 |  |  |
| **Bray-Curtis** | γ-diversity | 0.63 | 0.0042 | 152.35 | <0.001 | 0.72 | 25379.46 |
|  | sample size | -0.56 | 0.0042 | -135.6 | <0.001 |  |  |
| **Jaccard-Chao** | γ-diversity | 0.31 | 0.0067 | 45.88 | <0.001 | 0.26 | 41039.71 |
|  | sample size | -0.41 | 0.0067 | -60.61 | <0.001 |  |  |
| **normalized** | γ-diversity | 0.63 | 0.0038 | 165.51 | <0.001 | 0.77 | 22494.81 |
| **divergence** | sample size | -0.61 | 0.0038 | -159.6 | <0.001 |  |  |

**Table S2.**
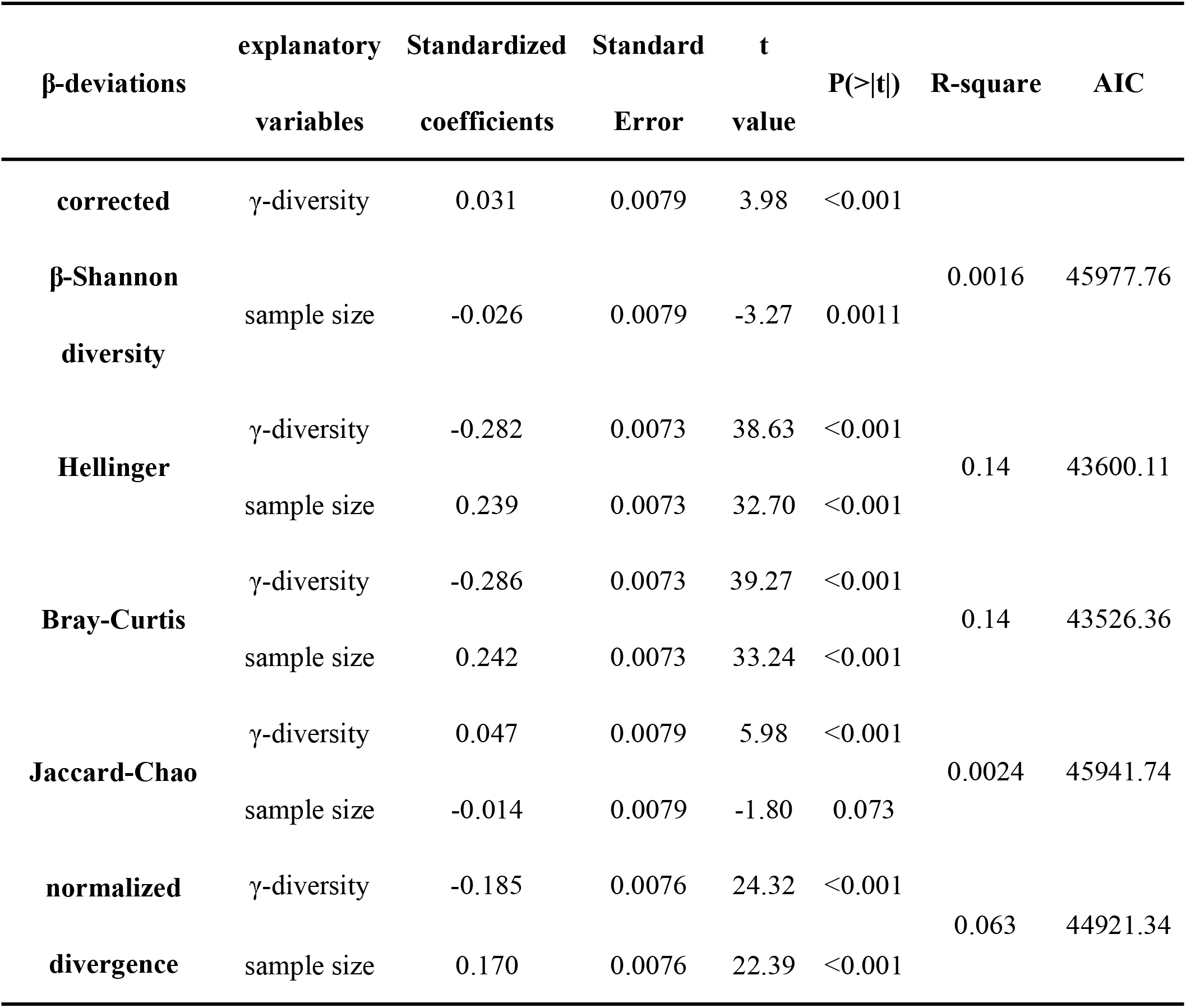
The sensitivity of different β-deviations (the deviation between the observed and null-expected β-diversity) to γ-diversity and sample size based on simulated communities. In each model, explanatory variables include γ-diversity and sample size. The standardized coefficients are standardized regression coefficients of multiple linear regression models.

**Table S3.** The sensitivity of β-metrics to γ-diversity and sample size based on Gentry’s global forest dataset. The standardized coefficients are standardized regression coefficients of multiple linear regression models.

| β-metrics | explanatory variables | Standardized coefficients | Standard Error | t value | P(>\|t\|) | R-square | AIC |
| --- | --- | --- | --- | --- | --- | --- | --- |
| **corrected** | γ-diversity | 0.44 | 0.079 | 5.66 | <0.001 | | |
| **β-Shannon diversity** | sample size | -0.094 | 0.079 | -1.20 | 0.23 | 0.16 | 531.53 |
| **Hellinger** | γ-diversity | 1.00 | 0.043 | 23.00 | <0.001 | | |
|  | sample size | -0.34 | 0.043 | -7.74 | <0.001 | 0.74 | 298.28 |
| **Bray-Curtis** | γ-diversity | 0.96 | 0.049 | 19.55 | <0.001 | | |
|  | sample size | -0.34 | 0.049 | -7.00 | <0.001 | 0.67 | 345.05 |
| **Jaccard-Chao** | γ-diversity | 0.95 | 0.052 | 18.15 | <0.001 | | |
|  | sample size | -0.52 | 0.052 | -9.88 | <0.001 | 0.63 | 370.60 |
| **normalized** | γ-diversity | -0.36 | 0.063 | -5.71 | <0.001 | | |
| **divergence** | sample size | -0.41 | 0.063 | -6.60 | <0.001 | 0.46 | 443.35 |

**Table S4.**
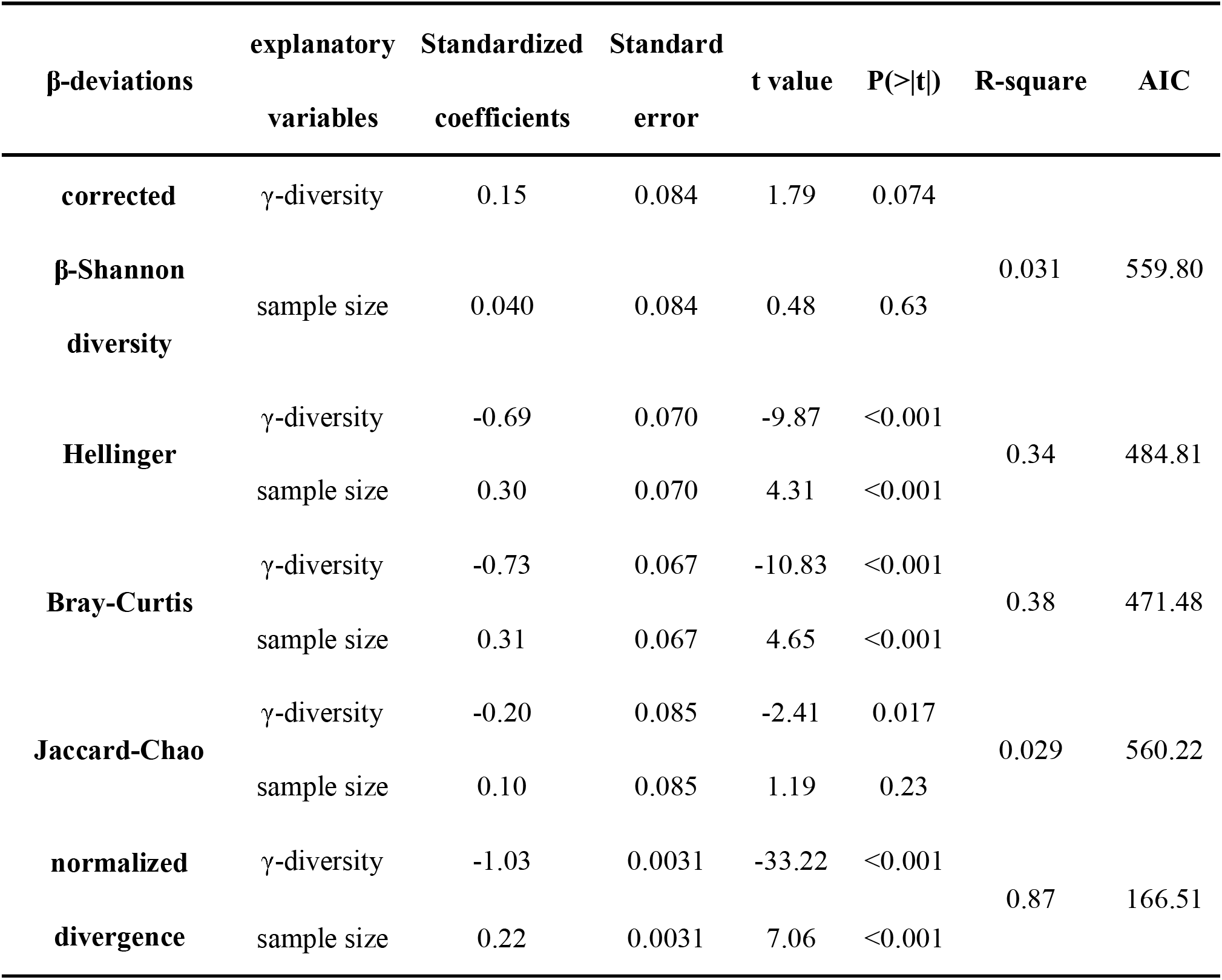
The sensitivity of different β-deviations (the deviation between the observed and null-expected β-diversity) to γ-diversity and sample size based on Gentry’s global forest dataset. The standardized coefficients are standardized regression coefficients of multiple linear regression models.

**Figure S1.**
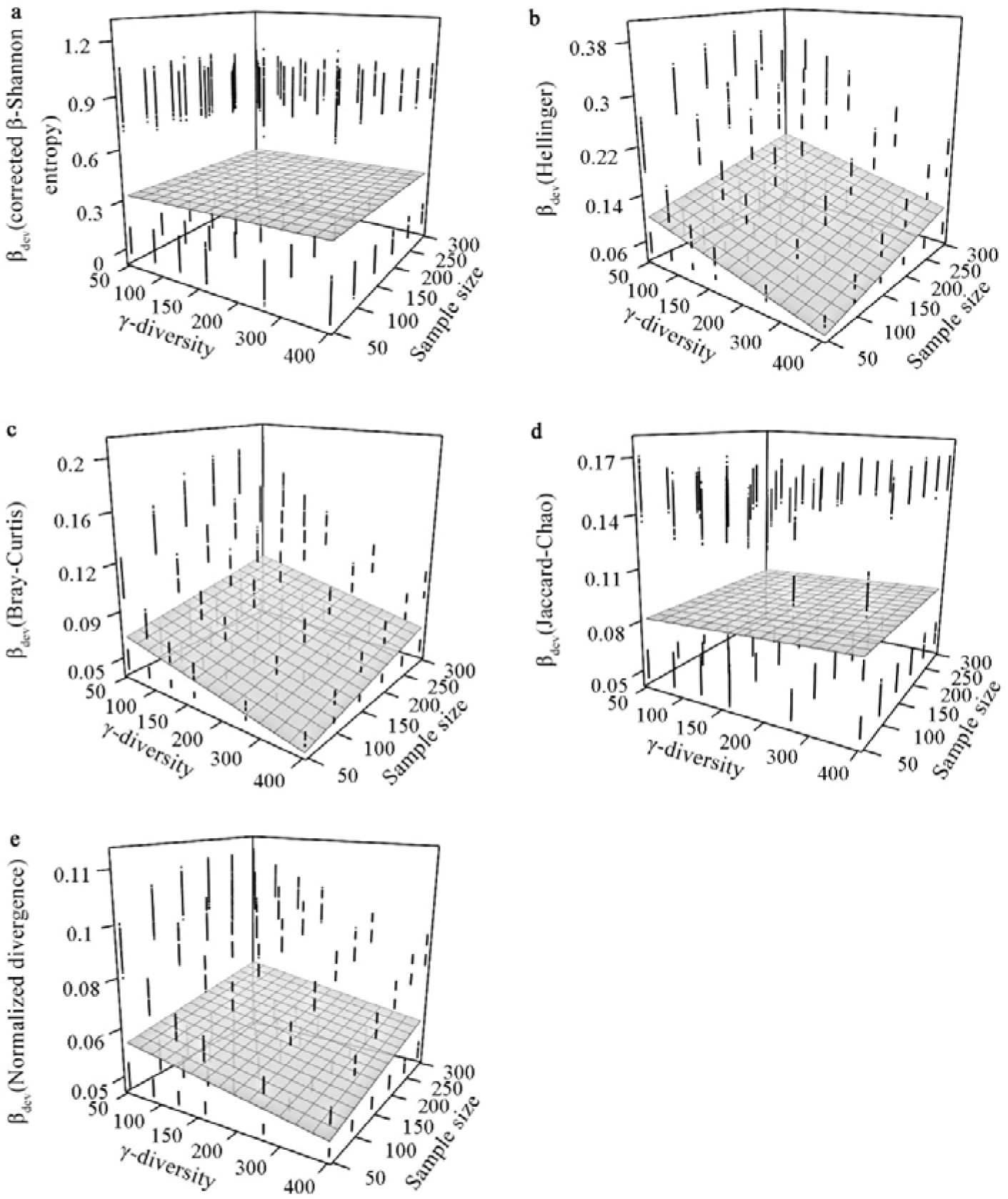
The sensitivity of β-deviation (β_*dev*_) to γ-diversity and sample size. β-diversity was measured by a) corrected β-Shannon diversity, b) Hellinger, c) Bray-Curtis, d) Jaccard-Chao, and e) the normalized divergence indices. In each panel, the surface of γ-diversity and sample size was fitted using multiple linear regression (detailed model parameters given in *Table S2*).

**Figure S2.**
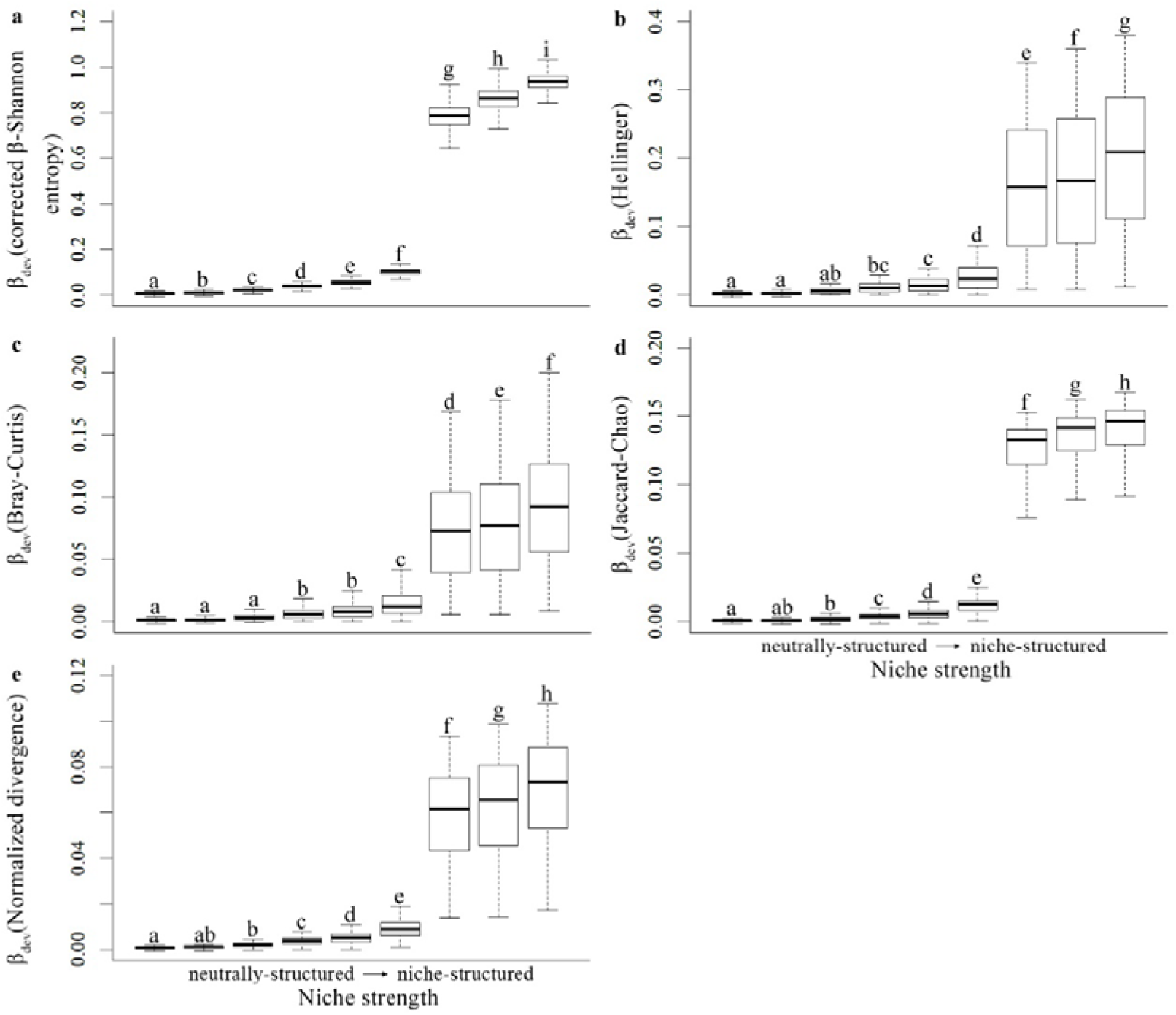
The variation of β-deviations (β_*dev*_) along a niche strength gradient. From left to right, metacommunities varied from stochastically to niche-structured; a) corrected β-Shannon diversity, b) Hellinger, c) Bray-Curtis, d) Jaccard-Chao, and e) the normalized divergence indices. The different letters annotated in each panel indicate significantly different mean values of different ecological scenarios.

## Supporting information

## Appendix 1. Five abundance-based β-metrics and their properties

## 1) *Corrected* β*-Shannon diversity*(Jost 2007)

β-Shannon diversity partitions γ-diversity into mathematically independent α- and β-diversity components as follows:

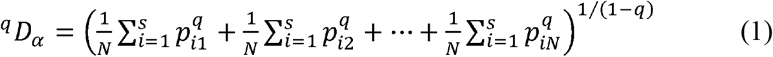

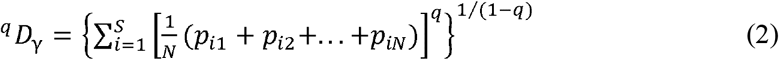

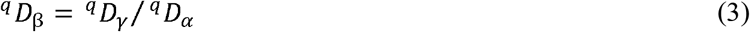

where ^*q*^*D*_*α*_,^*q*^*D*_*β*_ and ^*q*^*D*_*γ*_ are the *q*-th order α, β, and γ diversity, respectively; *p_i_* is the proportional abundance of species *I*; *S* and *N* are the total number of species and the total number of local communities (or plots), respectively, in the regional community. β-Shannon diversity is a standard abundance-sensitive measure that weights species in proportion to abundance according to diversity order *q*; when q = 0, *D* is equivalent to species richness. For increasing *q*, abundant species are given progressively more weight. Here we took *q* = 1 to weight all species by their abundance, without favoring either common or rare species. β-Shannon diversity quantifies β-diversity as the effective number of equally large and completely distinct communities, ranging from 1 to N (N is the number of local communities) (Jost 2007; Tuomisto 2010). We also incorporated undersampling correction methods with β-Shannon diversity for correcting γ-dependence (Chao *et al.* 2013; 2014).

## 2) *Normalized divergence* (Chao and Chiu 2016)

The normalized divergence unifies two major measures of β-diversity: the total variance of community species abundance matrix and diversity decomposition. The formulae are as follows:

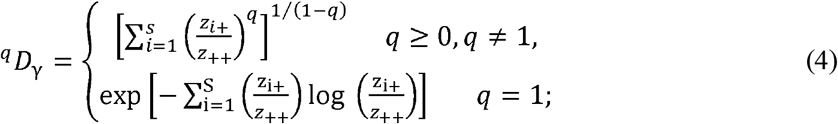

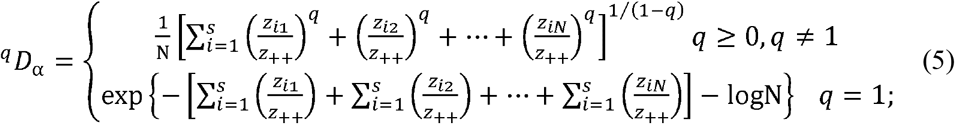

where z_i_ is the abundance of the *i*-th species in a local community, z_i+_ and z_++_ are the total abundances of the *i*-th species and of all species, respectively, in the regional community (Chao and Chiu 2016). Normalized divergence is calculated as the ratio of ^*q*^*D*_*γ*_ (equation 8) and ^*q*^*D*_*α*_ (equation 9). Focusing on abundance-based measures, ated as the ratio we took the value of diversity order *q* =1 for equal weight of rare and common species in this study, which is identical to β-Shannon diversity given true species richness and species abundance (Chao *et al.* 2019). Theoretically, normalized divergence uses a novel normalization method to transform the maximum value for completely distinct communities to 1, and removes these mathematical constraints by α, γ, and total abundance (Chao and Chiu 2016). However, it remains untested whether normalized divergence is robust to undersampling bias in empirical studies.

## 3) *Jaccard-Chao* (Chao *et al.*, 2006)

Jaccard-Chao, *D_J_*, is an abundance-based version of the classic Jaccard index. The equation for *D_J_* takes the form:

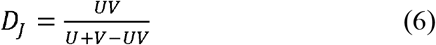

where *U* and *V* are the total relative abundances of individuals belonging to the shared species between two communities in community 1 and 2, respectively. Importantly, this index is mathematically independent of γ-diversity (Property 7 and 10 in Table 2 of Legendre and De Caceres 2013). Moreover, Jaccard-Chao corrects undersampling by incorporating undetected species into *U*, *V*—significantly reducing undersampling bias (Chao *et al.* 2005) (See more details regarding undersampling correction method for Jaccard-Chao in Appendix 2).

## 4) *Bray-Curtis* (Bray and Curtis 1957) *and Hellinger* (Rao 1995; Chao et al. 2015)

The abundance-based Hellinger (*D_H_*) and Bray-Curtis (*D_BC_*) indices are widely used pairwise dissimilarity measures of β-diversity. They are calculated as:

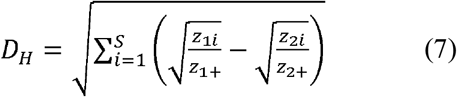

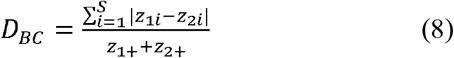

where z_1i_ and z_2i_ are the abundances of *i*-th species in community 1 and 2, while z_1+_ and z_2+_ are total abundances of community 1 and 2, respectively. The multivariate pairwise dissimilarity metrics, based on distance or dissimilarity matrices, quantify the average compositional dissimilarity across all pairs of samples in the study area. Moreover, both the Hellinger and the Bray-Curtis indices have the desirable property of being mathematically unrelated to γ-diversity (Table 6.2 in Jost *et al.* 2011; Property 7 and 10 in Table 2 of Legendre and De Caceres 2013). However, no undersampling correction methods are incorporated into either of the two metrics. We were especially interested in comparing β-diversity across multiple communities. Thus, we transformed the pairwise dissimilarity matrices quantified by Jaccard-Chao, Hellinger, and Bray-Curtis distance matrices into the total variance of community compositional heterogeneity as:

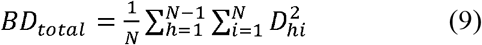

where *D_hi_* is the pairwise dissimilarity value of *i-*th and *h*-th position in the subdiagonal dissimilarity matrix (Legendre and De Caceres 2013). The total variance of the transformed data matrix is mathematically unrelated to γ-diversity and total abundance (Legendre and De Caceres 2013; Chao and Chiu 2016).

## Appendix 2. Undersampling correction methods and the null model approach

## Undersampling correction method for β-Shannon diversity

Chao *et al*. (2013; 2014) extended the species accumulation curve (Colwell *et al.* 2012) to the diversity accumulation curve, which corrects for the γ-dependence of β-diversity by asymptotically estimating the true α- and γ-Shannon diversity 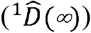 of samples in a region. The 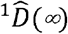 (*q*=1, *q* is the diversity order) is calculated as:

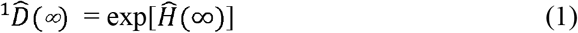

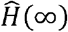 is a nearly unbiased estimator of Shannon entropy (Chao *et al.* 2013):

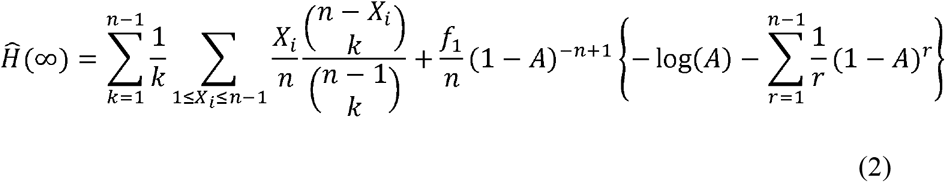

where *X_i_* is the species frequency of species *i*, *k* is the size of a random sample from the observed community, *f*_1_. is the number of singletons (i.e., species represented by only one individual in the observed sample), and *f*_2_ is the number of doubletons (i.e., species represented by only two individuals in the observed sample). *A* is the estimated mean relative frequency of the singletons in the sample:

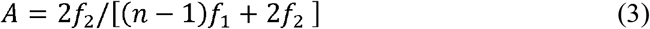

## Undersampling correction method for Jaccard-Chao

Chao *et al.* (2005) also proposed the undersampling-corrected Jaccard index (Jaccard-Chao), which estimates the relative abundances of undetected shared species.

The equation for Jaccard-Chao takes the form:

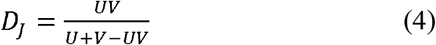

where *U* and *V* are the total relative abundances of individuals belonging to the shared species between two communities in community 1 and 2, respectively. The Jaccard-Chao index seeks to correct undersampling by incorporating undetected species into *U* and *V* (Chao *et al.* 2005). Specifically, the adjustment of *U* and *V* are:

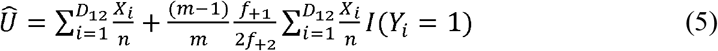

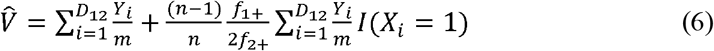

where *X_i_* and *Y_i_* are the numbers of individuals for the shared species *i* in communities 1 and 2, and *m* and *n* are the total abundances of all species in communities 1 and 2. *D_12_* is the number of shared species that are observed in both communities; *f*_+1_ and *f*_+2_ are the observed numbers of shared species that are singletons and and doubletons in community 2, while *f*_1+_ and *f*_2+_ are the observed numbers of shared species that are singletons and doubletons in community 1. Finally, *I(Y_i_=1)* is an indicator function, such that *I* = 1 when the shared species *i* is a singleton species in community 2 but may have any abundance greater than 0 in community 1, and *I* = 0 when the shared species *i* is not a singleton species in community 2; it also works for *I(X_i_=1)*.

## Randomization null model and β-deviation

The individual-based randomization null model approach simulates the effect of γ-dependence on β-diversity by randomly shuffling individual subplots of each plot (or among communities of each metacommunity in simulated data), while preserving the observed γ-diversity, the relative abundance of each species, and the number of stems in each subplot (Chase and Myers 2011; Kraft *et al.* 2011). We used 999 randomizations for each analysis. This method results in a β-deviation, indicating the difference in magnitude between the observed and expected β-diversity values (Chase and Myers 2011; Kraft *et al.* 2011). β-deviation values close to zero can be interpreted as stochastic assembly of communities, indicating the absence of deterministic processes, while those deviating from zero indicate that deterministic processes—such as habitat filtering or competitive interactions—cause patches to be more dissimilar than expected by chance (Chase and Myers 2011; Kraft *et al.* 2011).

## Appendix 3. Simulating metacommunities from strongly deterministic to relatively stochastic structure

To examine the robustness of β-metrics to γ-dependence in scenarios with strongly deterministic to relatively stochastic structures, we used a niche-based simulation model to generate the simulated metacommunities. The simulation models were constructed as follows:

1. We created a landscape of contiguous 20 × 20 subplots, in which each subplot represents a local community with adult trees selected from the species pool.
2. We generated two environmental variables (E) using an unconditional geo-statistical model, which was simply an application of the general Monte Carlo technique whereby values were generated by a particular variogram without constraints of real data (Webster and Oliver 2007). This technique allowed us to generate two environmental variables with known autocorrelation. That is, two arbitrary spherical semivariogram models with different ranges of autocorrelation, sill (sampled variance of a simulated environmental variable), and nugget (the local variation occurring at scales finer than the sampling interval) components to represent two environmental heterogeneity levels (Webster and Oliver 2007). The simulation of the environmental landscape was implemented using the R package ‘gstat’ (Pebesma 2004).
3. We created multiple simulation scenarios with the combination of different levels of γ-diversity, sample size, niche breadth (i.e., the environmental range that a particular species can tolerate), and niche position (i.e., the optimal environmental condition for a particular species) (Devictor *et al.* 2010). Subplots contained combinations of six levels of γ-diversity (50, 100, 150, 200, 300, and 400 species) and sample size (50, 100, 150, 200, 250, and 300 individuals). Recruitment dynamics were modeled as a lottery process, with species-specific survival probabilities determined based on niche breadth and niche position. By changing the combination of niche breadth and niche position, we simulated multiple metacomunities that varied in the degree to which deterministic versus stochastic processes were relatively important in response to an environmental gradient. In each scenario, all species were assigned with the same level of niche breadth but different niche positions (Tucker *et al.* 2016). Specifically, niche breadth (σ) was set equal for all species at three arbitrary levels (narrow: σ = 0.1, medium: σ =0.3, and wide: σ =0.5) for each species, where higher values confer larger ranges of species’ requirements in terms of habitat conditions (Gravel *et al.* 2006). Each species had a unique niche optimum (μ) and niche position, which were set to follow a β distribution that was symmetrically distributed around the mean habitat of the simulated metacommunity (Gravel *et al.* 2006). The β distribution is parameterized by two positive shape parameters, which control the shape of the distribution (skewness and kurtosis), denoted by α and β (not to be confused with α- and β-diversity) (Forbes *et al.* 2011). We set α and β to be equal to maintain symmetric distributions of niche positions. Like niche breadth, the aggregation of niche positions along the environmental axis was set at three levels by varying the kurtosis of the beta distribution (uniform aggregation: 2, moderate aggregation: 6, and high aggregation: 10) of shape parameters (α and β), correspondingly. This resulted in species distributed around the mean environmental habitats of a community with uniform, moderate, and high aggregation. For each scenario, we simulated 50 replicates.
4. In the simple niche-based competition model, we assumed that each species could reach all suitable subplots, and the interspecific inequality of competitive ability between species was mainly determined by the survival probabilities along the environmental gradient (Gravel *et al.* 2006). The survival rate of species *i* in each subplot was:

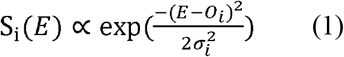

where *E* is the environmental value in each subplot, *o_i_* is the niche position of species *i*, and σ_*i*_ is the fundamental niche breadth of species *i*. By changing the combination of niche breadth and niche position, we simulated multiple metacomunities that varied in the degree to which deterministic versus stochastic processes influenced the responses to an environmental gradient. Metacommunities derived from deterministic processes arose when species were evenly distributed (i.e., uniform niche position) around the mean habitat condition and had a narrow niche breadth. Under this scenario, species had the largest interspecific differences in survival rate within each habitat, and competitive exclusion was strong. In contrast, stochastically assembled metacommunities arose when species were highly aggregated (i.e., aggregated niche positions) around the mean habitat conditions, and had a wide niche breadth. Under this scenario, all species had roughly the same competitive capability and survival rate across habitats (Gravel *et al.*, 2006).

